# MiMSI - a deep multiple instance learning framework improves microsatellite instability detection from tumor next-generation sequencing

**DOI:** 10.1101/2020.09.16.299925

**Authors:** John Ziegler, Jaclyn F. Hechtman, Ryan Ptashkin, Gowtham Jayakumaran, Sumit Middha, Shweta S. Chavan, Chad Vanderbilt, Deborah DeLair, Jacklyn Casanova, Jinru Shia, Nicole DeGroat, Ryma Benayed, Marc Ladanyi, Michael F. Berger, Thomas J. Fuchs, Ahmet Zehir

## Abstract

Microsatellite instability (MSI) is a critical phenotype of cancer genomes and an FDA-recognized biomarker that can guide treatment with immune checkpoint inhibitors. Recent work has demonstrated that next-generation sequencing data can be used to identify samples with MSI-high phenotype. However, low tumor purity, as frequently observed in routine clinical samples, poses a challenge to the sensitivity of existing algorithms. To overcome this critical issue, we developed MiMSI, an MSI classifier based on deep neural networks and trained using a dataset that included low tumor purity MSI cases in a multiple instance learning framework. On a challenging yet representative set of cases, MiMSI showed higher sensitivity (0.940) and auROC (0.988) than MSISensor(sensitivity: 0.57; auROC: 0.911), an open-source software previously validated for clinical use at our institution using MSK-IMPACT large panel targeted NGS data.

## Introduction

Microsatellite instability (MSI) is the phenotypic measure of deficiencies in the DNA mismatch repair (MMR) machinery, resulting in varying lengths of deletions and insertions in microsatellites. Microsatellites are short, tandemly repeated DNA sequences, and in patients with an MSI-high (MSI-H) phenotype, these regions exhibit significant errors due to replication slippage. The Food and Drug Administration (FDA) has approved pembrolizumab, an immune checkpoint inhibitor, for patients with MSI-H and/ or MMR-deficient (MMR-D) cancers of any histology ^1,2^. Thus, reliable and robust testing strategies for MSI/MMR status are critical for clinical management of patients with metastatic cancer. Traditional tools that allow screening of MSI status include MSI polymerase chain reaction (PCR) and/or MMR immunohistochemistry (IHC) testing but their application for pan-cancer screening including cancer types with a much lower prevalence of MSI than colorectal and endometrial cancer have raised concerns regarding cost-effectiveness and optimal resource and tissue utilization. Given that clinical comprehensive genomic profiling using targeted NGS panels is now being used more widely to inform treatment decisions in patients with advanced solid cancers, the advantages of also extracting MSI status from these data are apparent. We recently validated and implemented MSISensor as a way of identifying MSI status in patients who are undergoing next-generation sequencing (NGS) testing at Memorial Sloan Kettering Cancer Center (MSKCC) using MSK-IMPACT, an FDA-authorized targeted tumor sequencing panel ^3–5^. While this enabled a comprehensive and prospective MSI analysis across a wide array of tumor types, certain features of the clinical samples and the algorithm prevented us from reliably identifying all MSI-H patients. Because MSISensor calculates a distribution of the number of deletions in a given microsatellite region and compares the tumor and the matched normal distributions, samples with low tumor purity may lead to false negatives. Further we observed samples with low sequence coverage also suffer from false negative results. Finally, the presence of an ‘indeterminate’ category (MSISensor scores between 3 & 10, 3.8% of all samples tested), leads to complexities in patient management and the need for orthogonal testing with use of additional tissue resources.

The utility of supervised deep learning in classifying genomic data and results has been demonstrated by a number of somatic and germline variant callers based on deep learning methods ^6–8^. These prior genomic applications of deep learning methodologies have relied on labelled training data and are thus fully supervised. This existence of a true label for every datapoint makes learning in a fully-supervised manner computationally feasible. For MSI classification, the ground truth label is not at the individual genomic location as with the labels for individual variants. Rather, the label is for the entire sample and is based on a variety of testing modalities as previously mentioned. Here, we describe a new computational tool for accurately classifying MSI status using NGS data, called MiMSI (**M**ultiple-**i**nstance **MSI**, pronounced “My-MSI”). Our method utilizes a deep multiple instance learning model rather than traditional statistical modeling methods to achieve greater sensitivity while retaining specificity^9^.

## Results

### Model development

We utilized multiple instance learning (MIL), a weakly supervised machine learning methodology to develop an MSI classifier. With MIL, a broad label is applied to a collection of multiple individual data points, called instances, rather than to each instance individually ^10^. This problem formalization is evident in many machine learning applications in medicine and has since been combined with deep convolutional methods to form highly performant models ^11–13^. In formulating the MSI classification as a MIL problem, we consider each microsatellite region in a patient sample to be an instance and each patient sample to be our ‘bag’, or collection of instances. Ground truth MSI status exists at the bag level due to the orthogonal PCR and MMR-IHC testing performed for each case in our training and validation datasets. Therefore, we can compute the loss function and training accuracy for each sample without requiring the accurate classification of each individual microsatellite region. Optimizing the loss function across all the samples in our training dataset allows us to build a final model to predict the MSI status of subsequent test samples.

The end-to-end classification pipeline (**Figure 1a**) starts with converting the aligned NGS reads at each microsatellite site for both the tumor and matched normal sample into a vector representation. The model predicts a single probability that the sample is MSI by (i) calculating a feature representation for each microsatellite vector for a sample, (ii) averaging all microsatellite representations into a sample level embedding and (iii) utilizing a sigmoid classification layer on the preceding sample embedding to determine the final probability of MSI status (**Figure 1b)**. The first stage of the model is a deep convolutional neural network (CNN) designed to extract a feature embedding for each microsatellite vector in a given sample. The network (**eFigure 1**) is based on a ResNet architecture, employing residual connections after each pair of convolutional layers. These residual connections allow the network to combine both high-level and low-level features learned at varying levels in the network into one cohesive feature embedding at the final stages ^14^. The feature embeddings established by the network for each of the microsatellite loci are then averaged into a sample-level embedding. This final sample-level embedding is passed through the last stage of the network where we utilize a sigmoid classifier to predict the likelihood of MSI occurrence in a given sample.

**Figure 1.**
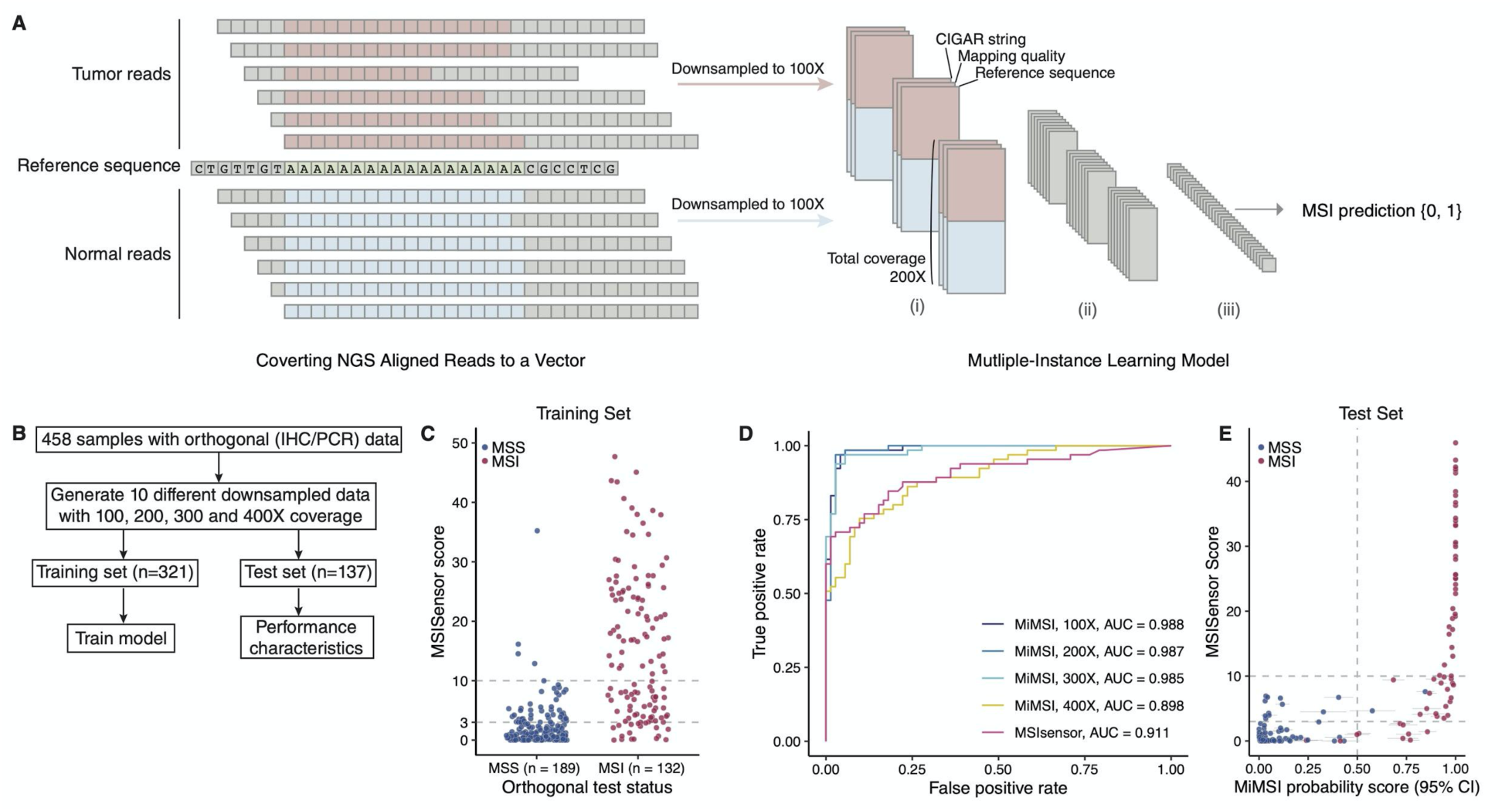
MiMSI model design and performance metrics. **A**. Schematic representation of converting sequencing reads in a given genomic region into vector representation. Reference sequence along with mapping qualities and CIGAR strings for each read is used in the vectorization after downsampling. The set of vectors for a given sample are passed through the model (see eFigure 1). **B**. Study cohort used for both training the model and testing the performance. **C**. Distribution of MSISensor scores for samples with orthogonal testing performed. **D**. Area under the receiver operator curve (auROC) analysis of the test cohort analyzed with MSISensor and MiMSI at 4 different downsampled coverage levels (100X, 200X, 300X, and 400X). **E**. MSISensor scores and MiMSI probabilities for the test cohort. Colors indicate the orthogonal test status.

### MiMSI Performance Compared to Orthogonal Testing

To train and validate the performance of MiMSI, we created a dataset, deliberately including cases where MSISensor had failed to identify the MSI status due to low tumor purity and coverage, and for which orthogonal MSI-PCR and MMR IHC data is available (n = 458, **Supplementary Table 1**). (**Figure 1c**). We trained the model using 321 samples with four different down-sampled coverage values when creating instance vectors: 100, 200, 300 and 400X, to normalize the coverage inequality between the tumor and the matched blood sample (**Figure 1d**, see **Methods**). Next, 137 previously unseen cases were used as a test set to compute the accuracy, sensitivity, specificity, and area under the receiver operator curve (auROC) metrics (**Figure 1e, Table 1**). MSISensor demonstrated a lower auROC (0.91), as we specifically selected cases that were challenging for MSISensor. Performance of MiMSI was highest with 100X and 200X models (auROC: 0.99 for both) and degraded with increased coverage. At higher coverages, down-sampling reduced the average number of loci used in training and testing (**eTable 1**). Both the 100X and 200X models demonstrated similar performance metrics, however, confidence intervals (CI) derived from performing random downsampling ten times showed 200X has less variation due to down-sampling (**eFigure 2**) therefore we used the 200X model for the rest of the analyses in this manuscript. Further, we used a score of 0.5 as a cut-off threshold to determine MSS vs MSI-H cases with MiMSI and classified any case for which the 95% CIs crossed the 0.5 boundary as MSI-indeterminate (MSI-ind).

**Table 1.** Performance metrics for MSISensor and MiMSI

| Method | Accuracy | auROC | Sensitivity | Specificity |
| --- | --- | --- | --- | --- |
| MSISensor | 0.805 | 0.911 | 0.571 | 0.985 |
| MiMSI 100X | 0.934 | 0.988 | 0.879 | 0.986 |
| MiMSI 200X | 0.963 | 0.987 | 0.940 | 0.986 |
| MiMSI 300X | 0.920 | 0.985 | 0.848 | 0.986 |
| MiMSI 400X | 0.788 | 0.898 | 0.697 | 0.873 |

**Figure 2.**
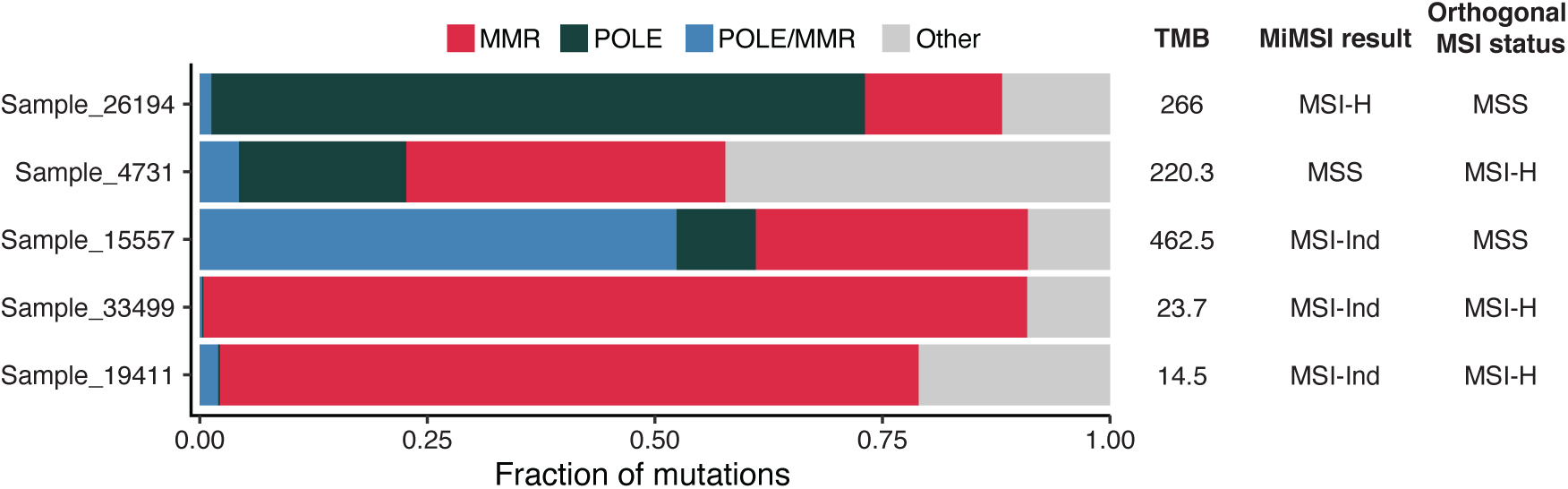
Mutational signature analysis for discrepant cases. Bar chart showing the fraction of mutations explained by a given mutation signature: mismatch-repair deficiency (MMR: Red); error-prone DNA Polymerase ε (POLE: grey) and concurrent MMR and POLE (blue). All other signatures are shown in light grey. TMB indicates the total number of mutations normalized by the genomic region covered by a given MSK-IMPACT panel version^5^. Due to low number of mutations (less than 14), we did not perform signature analysis for samples Sample_26600 and Sample_8865. Orthogonal testing is MMR IHC or MSI PCR.

Within the test dataset, we found a total of four indeterminate cases (two MSS and two MSI-H by orthogonal testing) and three discrepant cases (**Figure 2**). Amongst the discrepant cases, Sample_26194 was a colon adenocarcinoma with 303 somatic mutations detected and classified as MSI-H by MiMSI but MSS by orthogonal testing with IHC/PCR. Mutational signature analysis attributed 72% of the mutations to the deficiency of DNA polymerase-ε (POLE) resulting from an exonuclease domain mutation (V411L). 15% of the mutations were attributable to MMR deficiency and there was a nonsense mutation in *MSH2* (E580*), suggesting the possibility of a smaller clone with MMR phenotype that was either not clear during IHC review or MSH2 expression was retained ^15^. The two other discrepant cases were MSS by MiMSI score but MSI-H by orthogonal IHC/PCR. Sample_26600 was a uterine endometrioid carcinoma (UEC) with only three single nucleotide variants (SNVs) identified with a median variant allele frequency (VAF) of. 6.6%. The other sample, Sample_4731, was also a UEC with 224 SNVs and a median VAF of 7.7%. The low VAF of the mutations in these two samples suggest the tumor purity of these samples might be below the limit of detection.

Amongst MiMSI indeterminate and the orthogonally MSS cases, Sample_15557 was a UEC with over 500 mutations identified with only 5 indels. Mutation signature analysis showed 52% of the mutations were attributable to concurrent deficiencies in proficient DNA replication by POLE and MMR system (Figure 2) <u>^16^</u>. The other case, Sample_8865, was a uterine clear cell carcinoma with only 5 mutations identified. Amongst the orthogonally MSI-H cases, Sample_33499 was an endometrial cancer case with 27 mutations identified, with a median VAF of 5% suggesting a very low tumor purity. In this instance, mutation signature analysis showed 90% of the mutations contributing to MMR deficiency. Sample_19411 was also a UEC with 16 mutations with low tumor purity, where the mutations signature analysis showed 77% of the mutations contributing to MMR deficiency. In both instances, signature analysis of mutation contexts could help clarify the MiMSI indeterminate cases.

We further performed sample dilution experiments to determine the sensitivity of MiMSI to changes in tumor purity, and to compare its performance at low purity with MSISensor (**eTable 2**). We diluted a tumor DNA sample from an MSI-H case validated by MSI PCR with successive amounts of normal DNA from matched FFPE normal tissue and evaluated the sequencing data with both MSISensor and MiMSI. The MSISensor score decreased from its original value of 36.7 as tumor purity decreased, however it remained indicative of an MSI-H phenotype until the final dilution of 6%. At that point MSISensor lacked sufficient signal to score the case above our MSI-H threshold of 10. MiMSI, however, classified the sample as highly likely to be MSI-H, even at the lowest dilution point.

**Table 2.** Comparison of MiMSI and MSISensor results

|  |  | MSISensor |  |  |
| --- | --- | --- | --- | --- |
|  |  | MSS | MSI-Ind | MSI-H |
| MiMSI | MSS | 39,173 | 1,388 | 52 |
|  | MSI-Ind | 1,327 | 89 | 6 |
|  | MSI-H | 455 | 212 | 998 |

### Importance of number of microsatellite sites processed

Since vector data generated over the microsatellite sites are used for the prediction of MSI-H phenotype, we investigated how the number of loci used in analysis affects the outcomes. We randomly downsampled the microsatellite loci used in the test cohort and asked how MiMSI classifications change (**Figure 3**). As the number of loci used decreased, the confidence intervals of the samples analyzed increased, which led to an increase in the number of MSI-Ind cases. Rate of MSI-Ind cases increased from 2% (1,000 sites used) to 27% (100 sites used) demonstrating that the number of sites used is an important factor in classifying cases properly.

**Figure 3.**
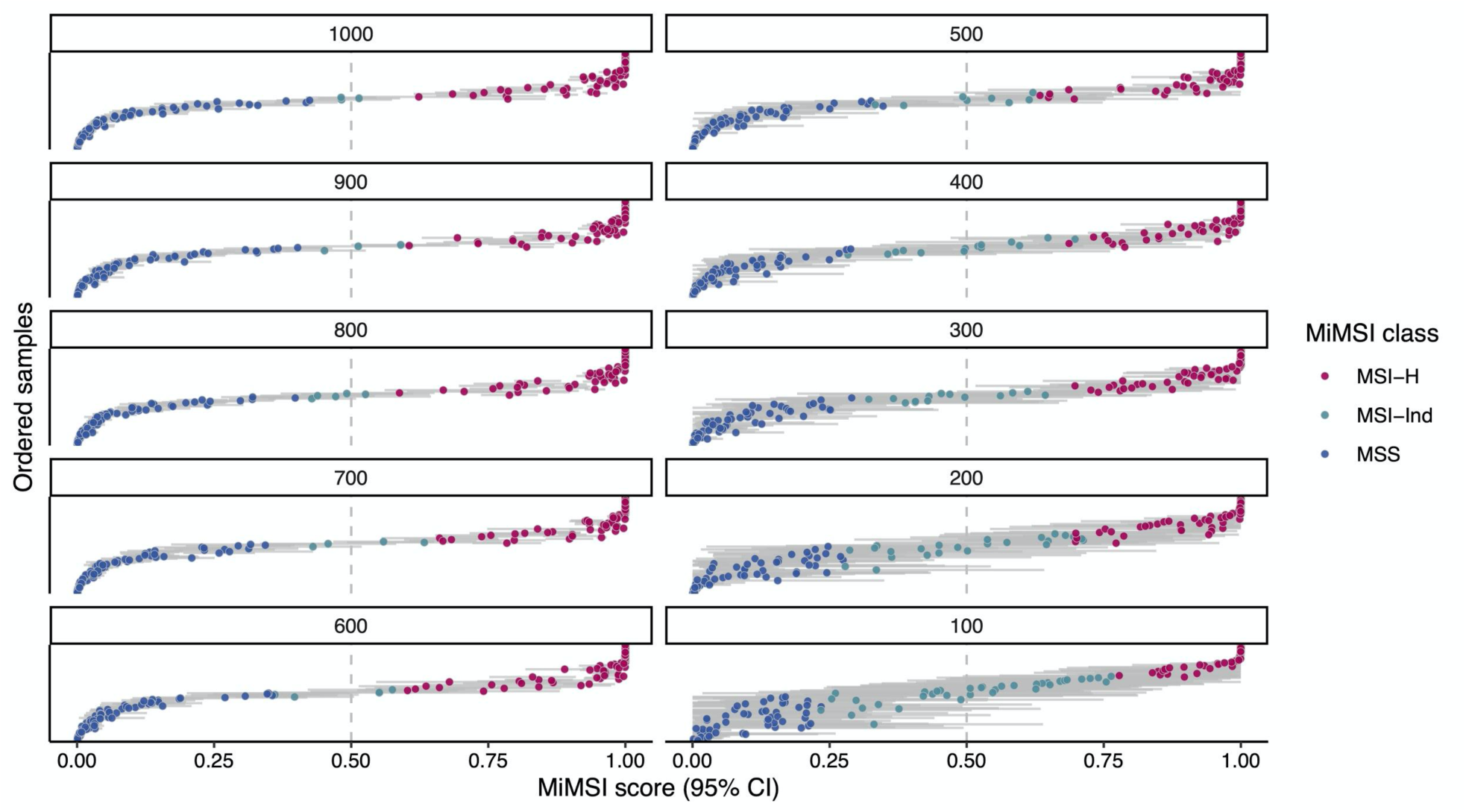
Downsampling microsatellite loci used in classification. MiMSI classification results after randomly downsampling microsatellite loci used for the classifier.

### Analysis of tumors without a matched control

While MiMSI was trained using a patient-matched normal control sample, there are instances where a matched control may not be available. In order to assess MiMSI’s ability to classify cases in a tumor-only mode, we reanalyzed the samples in the test cohort using an unrelated normal sample (**Figure 4a**). We observed that while the majority of the orthogonally MSS cases had scores < 0.5, orthogonally MSI-H cases had scores < 0.5 as well leading to false negative (FN) calls. We hypothesized that ethnicity differences between the tumors and the normal comparator used could lead to FNs. Therefore, we also tested a pooled blood control (an equimolar mixture of 10 blood samples) as comparator (**Figure 4b**). The results were similar to single unmatched normal comparator suggesting ethnicity may not be the reason for FNs. We noted that the majority of FNs in this setting were tumors of low tumor purity where MiMSI might be leveraging data from matched normal sample to increase its sensitivity. Therefore, we suggest users be cautious in interpreting the results and maybe want to apply different thresholds to reduce the false negative results.

**Figure 4.**
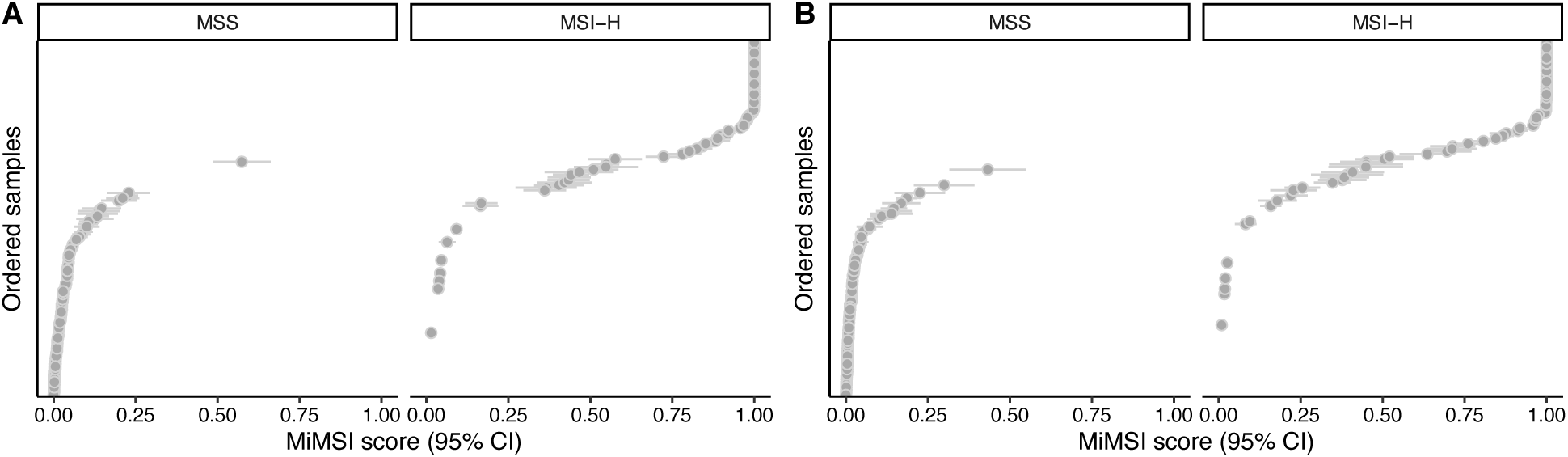
Tumor-only analysis of test cohort. Test cohort was analyzed using three different pooled normal samples.

### Comparison of MiMSI with MSISensor

After validating the results of MiMSI, we set out to investigate MMR-D phenotype across a large variety of cancer types (n = 86) by analyzing 44,724 tumor samples sequenced with the MSK-IMPACT assay. Comparison of MiMSI results with MSISensor showed overall 91% concordance for cases identified as MSS and MSI-H with both methods (**Figure 5, Table 2**). Moreover, MiMSI reduced the number of samples in the MSI-Ind category identified by MSISensor from 3.8% (n=1,689) to 3.2% (n=1,422).

**Figure 5.**
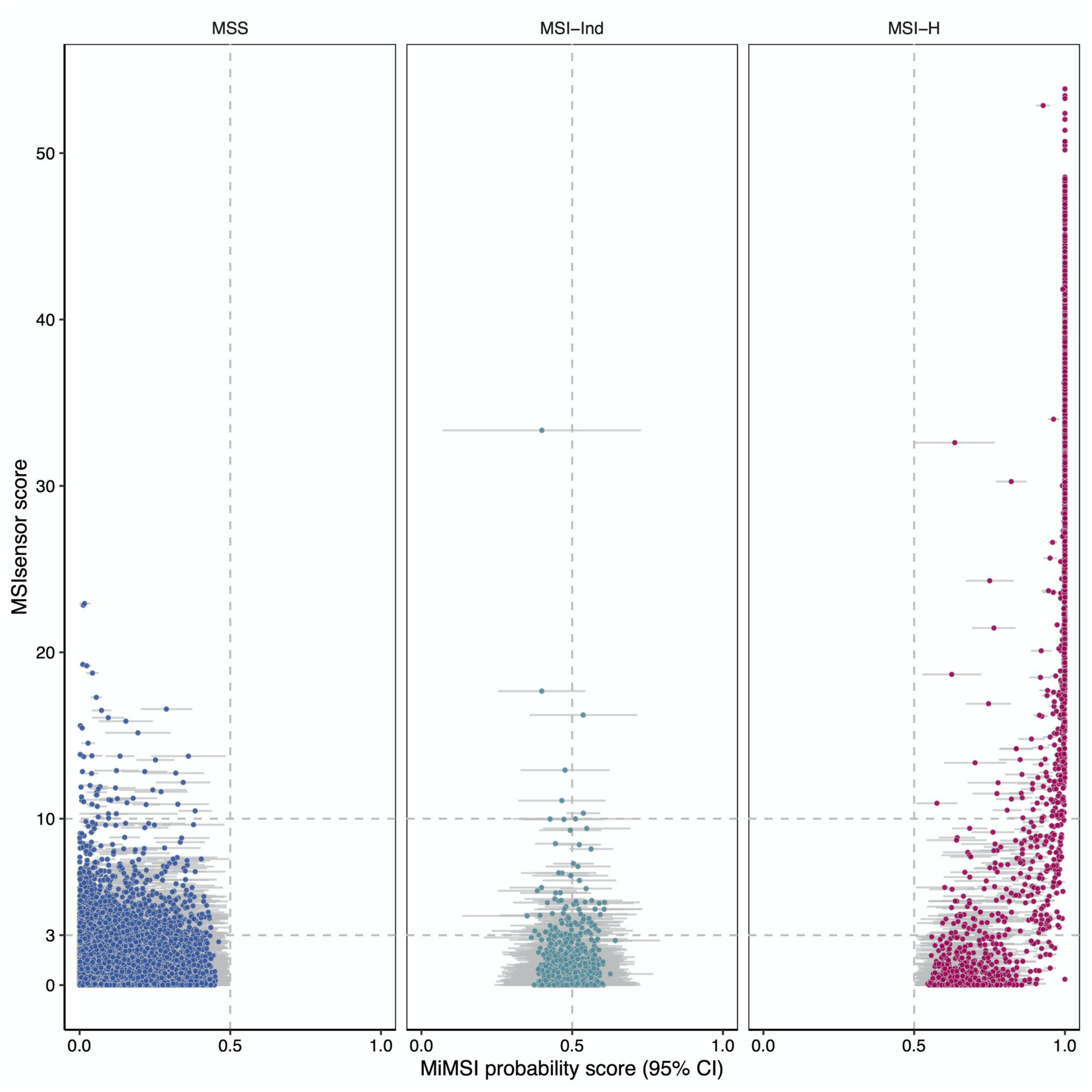
Global comparison of MiMSI results with MSISensor. Comparison of MSISensor scores and MiMSI probability scores (95% CI) for a cohort of 44,724 tumor samples

## Discussion

Here, we describe a novel algorithm for MSI classification caller which uses vectorized NGS reads at microsatellite loci and is extensible to multiple DNA capture types. Given the importance of correctly identifying the MSI status in clinical cohorts, and the fact that low tumor purity is common in formalin-fixed paraffin-embedded specimens, we believe MiMSI will be a valuable addition to clinical analysis pipelines for comprehensive genomic profiling.

Detecting MSI-H status is crucial for the management of cancer patients: it is predictive of Lynch syndrome regardless of the primary tumor as well as predictive of response to immune checkpoint inhibition^17^. However, current DNA based methods often have limited technical sensitivity in many MSI-H samples as these tumor samples typically show increased levels of tumor-infiltrating lymphocytes or a ‘Crohn’s-like’ lymphocytic reaction, which can sharply reduce tumor purity (i.e. the proportion of all cells in the biopsy that are tumor cells). To date, MMR IHC is the only tumor purity agnostic method to screen for MSI-H status, but it requires separate material which cannot be used for other predictive tests (such as sequencing). We show that MiMSI can be used concurrently with NGS assays performed on low tumor purity cases, decreasing the need for additional follow up IHC tests required with MSISensor. Further, we show MiMSI can detect MSI-H phenotype that occurs concurrently with other genomic lesions such as exonuclease domain mutations in *POLE*, where the MSI phenotype may not be clearly apparent. Finally, MMR IHC has a sensitivity of approximately 94% results can show false-retained MMR protein patterns with pathogenic missense mutations, removing an indication for treatment with immune checkpoint inhibition and points to the need for sensitive testing methods^15^.

The application of a deep learning-based classifier to NGS data allows us to infer MSI status with greater sensitivity than prior statistical methods, especially in situations where the sample has low tumor content or sequencing produces many low coverage regions. Since these traditional statistical methods rely on the difference in distributions between the tumor and normal sample they are sensitive to sequencing issues or sample quality issues. Our proposed model is trained utilizing a dataset composed of samples with varied levels of coverage, purity, and sequencing quality, resulting in a more resilient classifier. Additionally, the deep nature of our model means that it is able to classify against multiple levels of features derived from the aligned sequencing results, rather than just a set of hand-chosen features such as the number of deletions observed.

The results demonstrate that a deep multiple instance learning approach can be utilized to infer abnormalities in NGS data, even when that data is weakly labeled. Microsatellite instability is one such application; however, this method is applicable to other classification tasks as well. As NGS assays become more prevalent in clinical diagnostics methods like MiMSI allow us to identify MMR deficient and MSI-H cases during sequencing rather than as a separate IHC or PCR test. Furthermore, MiMSI can be utilized across many cancer types, raising the probability that additional MSI cases can be identified in tumor types where MSI testing isn’t typically performed or MMR deficiency isn’t commonly observed.

## Methods

### Clinical data generation

Prospective clinical sequencing data from 44,724 samples (40,064 patients) based on MSK-IMPACT panel ^4^ were used with MSKCC Institutional Review Board approval. MSISensor(v0.2) was used in the clinical analysis pipeline. MSISensor scores between 0 and 3 were considered MSS, between 3 and 10 were considered MSI-Ind and scores greater than were considered MSI-H ^3^. Based on our analytical validation, for samples with tumor content less than 20% MSI classification based on MSISensor is not reported.

### Converting NGS Data to Vectors

The intermediate file from which variant analysis is typically performed in NGS is the alignment file (BAM/SAM). The format of the alignment file is not conducive for implementation of deep learning algorithms involving a convolutional neural network (CNN) approach. Thus, we internally developed a program that efficiently converts the known microsatellite regions of the genome to three-dimensional vector representations of fixed sizes.

Each microsatellite region in the alignment file for both the tumor and normal sample is read utilizing pySAM, a python-based SAMtools reader. An individual vector for every site above a coverage threshold is generated by compiling a list of the aligned reads that completely span each given microsatellite region. Each aligned read is subsequently converted into a two-dimensional vector of size L ⨯ 3, where L is the length of the microsatellite region on the reference sequence and three is the number of data points extracted from the alignment file at each base. These three channels are the CIGAR string value, the mapping quality, and a binary integer representing the reverse strand mapping.

The generated single-read vectors are concatenated to form a three-dimensional region-level representation with dimensions *C* ⨯ *L* ⨯ *3*, where *C* is the required coverage at the interested region. In order to achieve the standard input size the CNN requires we downsample the coverage to the desired cutoff threshold for the first dimension. In our experiments we used coverage cutoffs of 50, 100, 150 and 200. The downsample process is random as to avoid bias from sorting of the alignment file in our model. Because the microsatellite regions vary in size and the model requires fixed dimensions, we introduced zeroes on the edges of the vectors for microsatellite regions that are smaller than 40 bp. This is referred to as zero-padding. This process results in *C* ⨯ *40* ⨯ *3* vectors for both the matched tumor and normal samples at each microsatellite locus. With the assumption that comparison between tumor and normal is necessary, we stacked the two vectors together, achieving a *2C x 40 x 3* vector for every microsatellite region in the patient’s genome. Due to coverage variability across various regions, the number of microsatellite vectors varies by patient. Our final input for training and testing the model is an *N* ⨯ *2C* ⨯ *40* ⨯ *3* vector for each patient sample, where *N* refers to the number of microsatellite vectors generated for the particular patient sample.

### MiMSI Model Architecture

MiMSI’s architecture (**eFigure 2**) can be separated into two main components. The first portion of the model creates an instance-level feature embedding for every microsatellite site, and the second calculates a bag-level embedding on which we calculate our final MSI prediction.

The instance-level model is based on ResNet-18, an 18-layer deep convolutional neural network that has performed well in various image-based classification tasks^18^. The original ResNet-18 architecture was adapted in order to accommodate the small size of our microsatellite vectors and to support a multiple instance learning approach. Compared to the 18-layer ResNet-18, MiMSI was built with 12 layers, and we downsampled the image twice over the course of the forward pass rather than at every residual connection. In each standard convolution layer a kernel size of 3 ⨯ 3 is utilized, with zero padding and a stride of 1. In the downsampling convolutions the kernel size, padding and stride are 3 ⨯ 3, 0, and 2 respectively. The stride is increased to 2 in order to downsample the vector by a factor of 2x.

The output of the final residual block in the instance-level model is fed into a fully connected linear layer to create an N ⨯ 512 vector representation of the sample, where N is the number of microsatellite loci. This collection of instance-level feature embeddings is further processed in the second portion of the model, which combines the instances into a bag-level score. We calculate the mean over the N microsatellite sites to form a bag-level embedding of 1 x 512. The final prediction of MSI status is calculated via a sigmoid layer on the preceding bag-level embedding. The final output of the model is a probability of MSI status, which we threshold at .5 in order to achieve our final binary classification of MSS vs MSI-H.

### Dataset

Our training and validation datasets consisted of 458 cases for which we had orthogonal MMR IHC or MSI PCR testing. These cases were a combination of cases from internal validation cohorts as well as 24 cases that were labeled as exceptionally difficult based on purity, coverage, or MMR status. The ground truth labels were determined via IHC or PCR testing for MMR-d and/or MSI status. MSI cases were given a ground truth label of one, while MSS cases were labeled zero. Our training dataset was a subset of 321 of the full 458 labeled cases. The training dataset contained a total of 189 MSS cases and 132 MSI-H cases. The remaining 137 cases were kept unseen from the model and used as our test dataset.

### Training and Validating the Model

The model is trained by minimizing the binary cross entropy loss function across the training dataset of 321 cases. The loss function is given by:

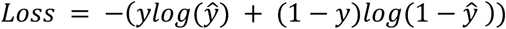

The model was optimized using the Adam algorithm with multiple learning rates and weight decay and the model with the highest accuracy on the testing set was chosen^18^. In our experiments we found the best performing learning rate and weight decay to be .0001 and .0005 respectively. The model was trained for 60 epochs, and initialized with random weights. The batch size was set at one, meaning that each epoch was one pass through every sample in the training dataset.

To validate our model we tested an unseen dataset of 137 cases, of which 72 were MSS and 65 were MSI-H, and we report our metrics against those held out samples. To compare performance against MSISensor we ran the tool against the same 137 cases utilizing the default parameters.

### Downsampling Effects on Accuracy

Evaluation on our held-out test set indicated that the 100x and 200x coverage models were the best performing models. However, given that MSK-IMPACT is a targeted, deep sequencing assay with average coverage close to 600x we conducted further analysis to quantify the effects of randomly downsampling reads down to these lower coverage levels before inputting the generated vectors into our model. Each case in the test set cohort was randomly downsampled to 100 and 200 reads 10 times each, giving us 10 replicates of each tumor/normal sample to input into the trained 100x and 200x models, respectively. Each replicate was evaluated using its corresponding model resulting in 10 predictions, which we used to create 95% confidence intervals for each of the 137 samples at both coverage levels. Comparing these confidence intervals, the 200x model empirically demonstrated more stable predictions (**eFigure 2**). The average length of a prediction’s confidence interval was .044 for the 200x model, compared to .06 for the 100x model. Also, less confidence intervals overlapped our prediction threshold of .5 when using the 200x model. This is particularly important for our application, since a confidence interval that overlaps our threshold effectively creates an “Indeterminate” case. Based on our initial goal of limiting indeterminate cases and to ensure the reproducibility of our predictions we opted to utilize 10 downsampled replicates at 200x with the higher coverage model to analyze our full cohort of approximately 45,000 MSK-IMPACT samples and to compare concordance with MSIsensor.

### Error, Sensitivity and Specificity

A case with an MSISensor score of more than 10 was considered MSI-H, while a case with a score between 3 and 10 was considered Indeterminate, and a score less than 3 was considered MSS. A case with a probability of MSI-H less than .5 by MiMSI was considered an MSS classification, while a score greater than or equal to .5 was considered MSI-H.

Error Rate (E), Sensitivity (TPR) and specificity (TNR) were calculated according to the following formulas:

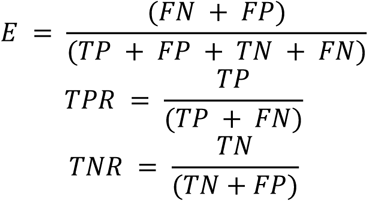

For calculating MSISensor accuracy we considered a case a true negative if MSISensor returned a score less than 3 for a case confirmed MSS by PCR/IHC, or a true positive if MSISensor returned a score greater than 10 for a case confirmed MSI-H by PCR/IHC. A case was considered a false negative if MSISensor classified a case Indeterminate or MSS and it was determined to be MSI-H by orthogonal testing, and a false positive if MSISensor classified a case MSI-H that was classified MSS by PCR/IHC. The Receiver Operating Characteristic (ROC) and Precision Recall plots were built utilizing the functionality in the open source scikit learn package, and we calculated the Area Under the Receiver Operating Characteristic (auROC) from this package as well.

### Concordance with MSISensor on the MSK-IMPACT Cohort

The MSK-IMPACT cases were analyzed with the default parameters of MSISensor and subsequently classified by the MiMSI model after training and validating on the dataset described above. We retained the same cutoffs for MSS, Indeterminant, and MSI-H for MSISensor cases as utilized in training and validating the model above. We defined a concordant case as one where (1) both MSISensor and MiMSI classified the sample as MSS or (2) both MiMSI and MSISensor classified the sample as MSI-H. Any other combination of classifications from MSISensor and MiMSI were treated as a discordant result.

### Mutational signature analysis

Mutational signatures were determined as described before ^6^

## Code Availability

The code developed and used in these experiments has been made freely available under version 3 of the GNU General Public License (GPLv3). The software along with the fully trained model is hosted at https://github.com/mskcc/mimsi

## Acknowledgements

We thank the members of Molecular Diagnostics Service and Fuchs Laboratory for their input. This study was funded by the National Cancer Institute under the MSK Cancer Center Support Grant/Core Grant (P30 CA008748).

## Extended Data and Figures

**Extended Table 1.** Average number of sites utilized for each coverage downsampling analysis by MiMSI in the testing cohort. Minimum and maximum numbers used are shown in parentheses

| Coverage | Avg # of sites (min-max) |
| --- | --- |
| 100X | 932 (424-1412) |
| 200X | 670 (190-1027) |
| 300X | 494 (74-816) |
| 400X | 363 (26-723) |

**Extended Table 2.**
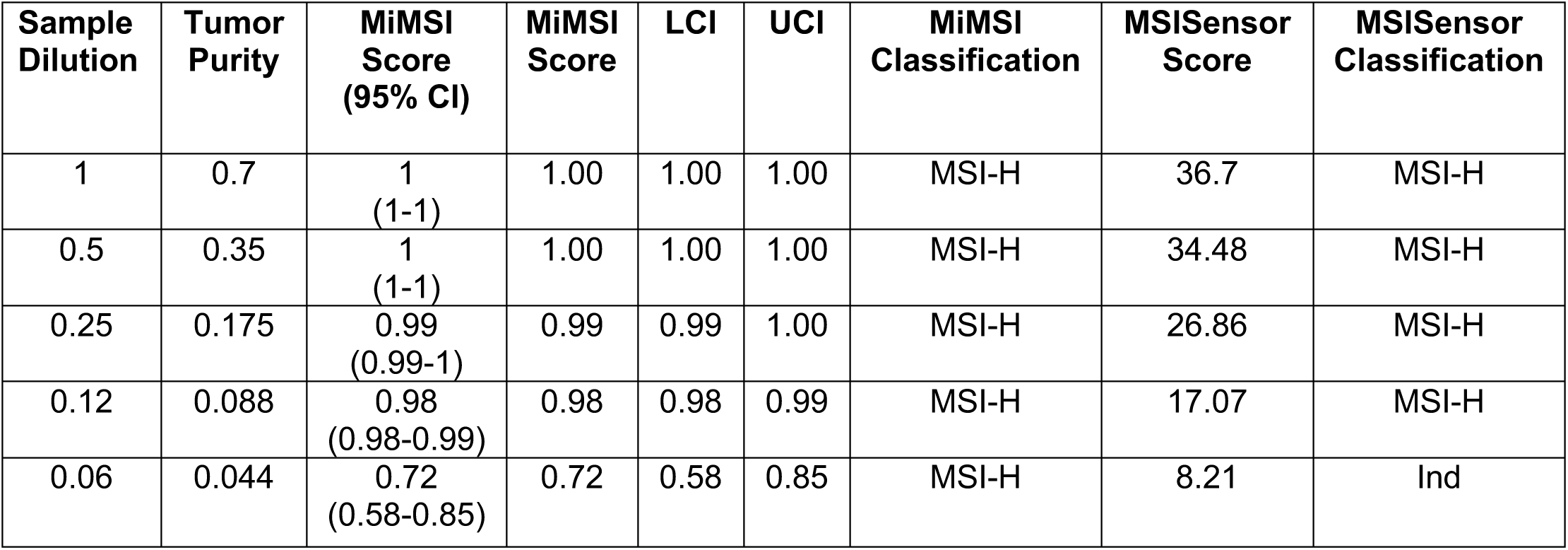
MiMSI and MSISensor analysis of dilution series. Tumor purity was calculated using FACETS.

**Extended Figure 1.**
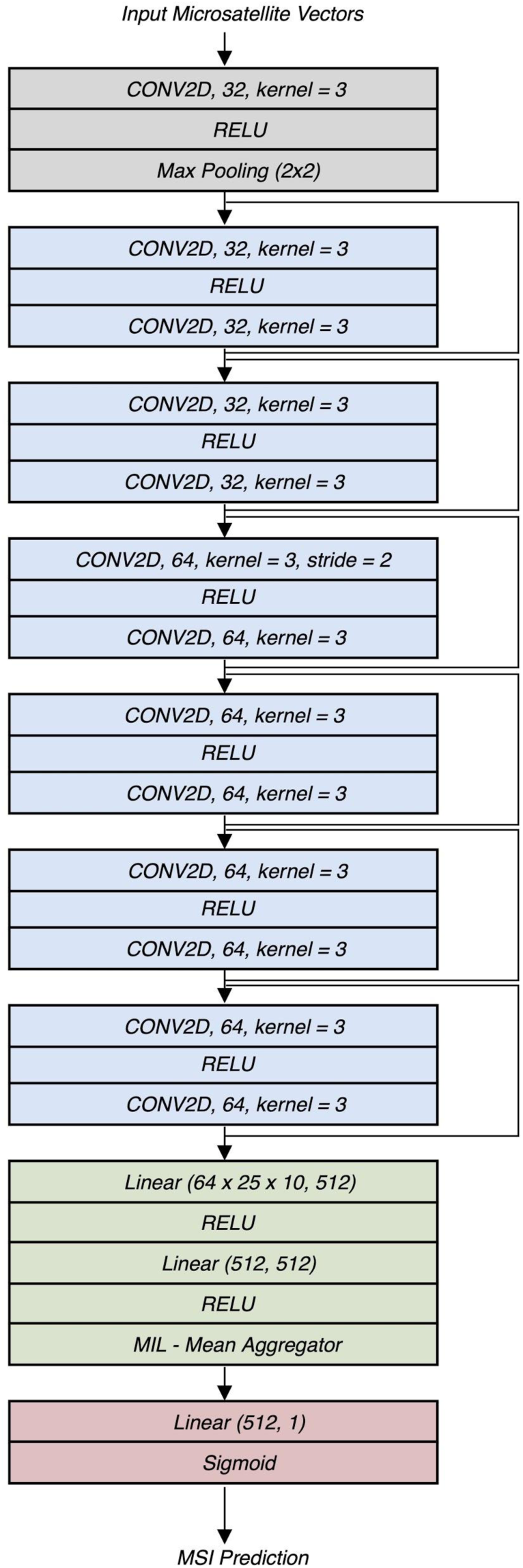

**Extended Figure 2.**
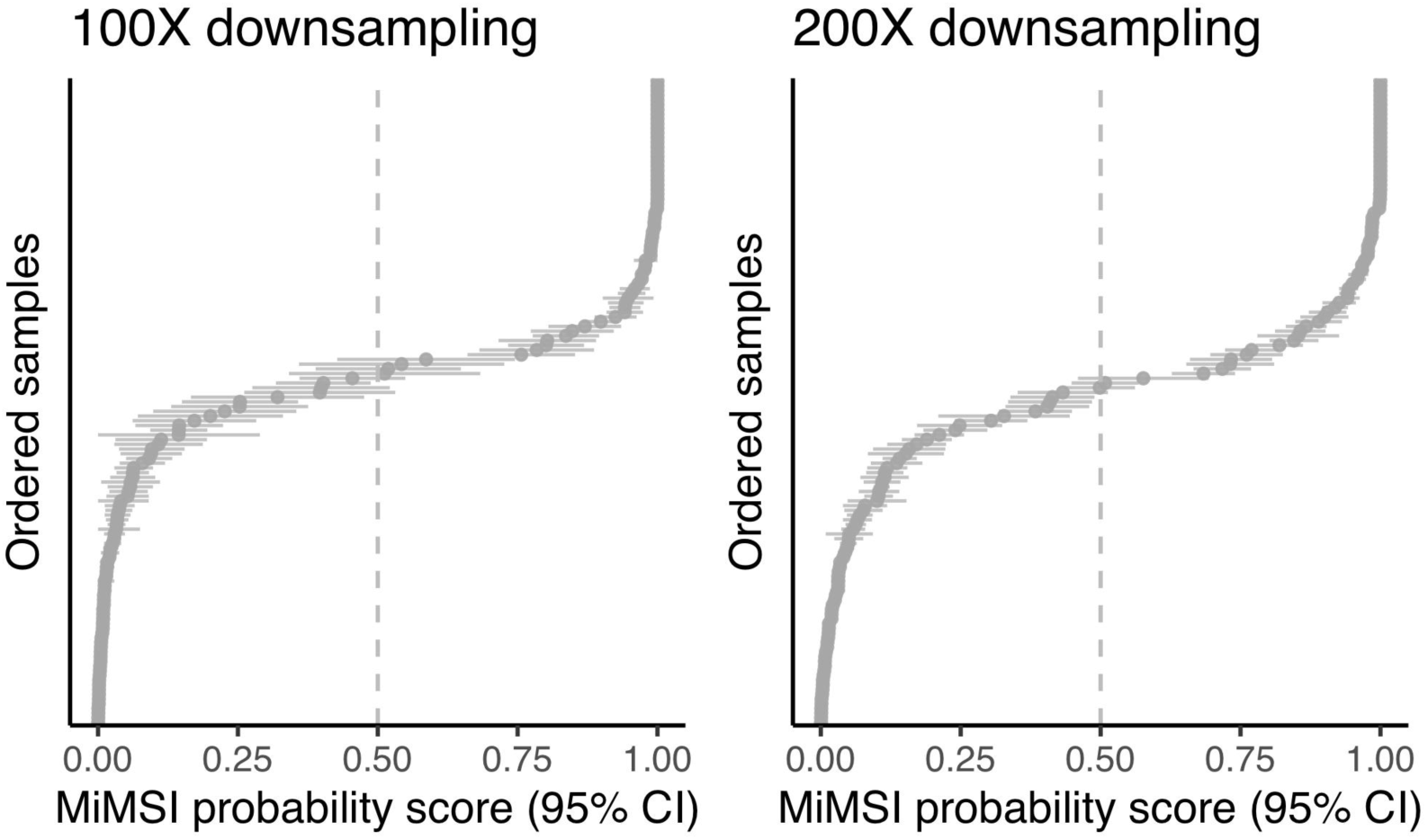
Range of MiMSI probability scores for each sample across 10 replicates with 100X (**A**) and 200X (**B**) downsampling

## Supplementary Data

**Supplementary Table 1**. Characteristics of the samples used as training and test datasets

